# Osmoregulation in freshwaters: Gene expression in the gills of a Neotropical cichlid in contrasting pH and ionic environments

**DOI:** 10.1101/489211

**Authors:** Stuart C. Willis, David E. Saenz, Gang Wang, Christopher M. Hollenbeck, Luiz A. Rocha, David S. Portnoy, James J. Cai, Kirk O. Winemiller

## Abstract

Freshwater habitats of the Neotropics exhibit a gradient from relatively neutral, ion-rich whitewater to acidic, ion-poor blackwater. Closely related species often show complementary distributions among ionic habitats, suggesting that adaptation to divergent osmoregulatory environments may be an important driver of Neotropical fish diversity. However, little is known about the evolutionary tradeoffs involved in ionoregulation across distinct freshwater environments. Here, we surveyed gill mRNA expression of *Cichla ocellaris* var. *monoculus*, a Neotropical cichlid, to examine cellular and physiological responses to experimental conditions mimicking whitewater and blackwater.

Gene ontology enrichment of expressed genes indicated that the gills were remodeled during both forms of environmental challenge, with changes biased towards the cellular membrane. We observed expression of signaling pathways from both the acute and extended response phases, including evidence that growth hormone (GH) may mediate osmoregulation in whitewater through paracrine expression of insulin-like growth factor I (IGF-I), but not through the GH receptor, which instead showed correlated up-regulation with the prolactin receptor and insulin-like growth factor II in blackwater.

Differential expression of genes related to paracellular tight junctions and transcellular ion transport showed responses similar to euryhaline fishes in fresh versus seawater, with some exceptions, suggesting that relaxed ion retention via the gills, possibly mediated by the GH/IGF-I axis, is a strong candidate for evolutionary modification in whitewater and blackwater endemic populations. In each osmoregulatory domain, we saw examples of contrasting differential expression of paralogs of genes that are single copy in most terrestrial vertebrates, indicating that adaption by fishes to diverse physicochemcial environments has capitalized on diversification of osmoregulatory gene families.

## Introduction

Gill surfaces of fishes must be thin in order to facilitate gas exchange, but this also promotes rapid gain or loss of ions and water with the environment – a functional tradeoff called the ‘osmo-respiratory compromise’ [1]. Management of osmotic and ionic stress is therefore a major biological challenge for fishes and has been estimated to account for 2-20% of resting metabolic rate [2]. A great deal has been learned about mechanisms involved in osmoregulation from studies of euryhaline fishes exposed to salinity gradients [3]; however, these fishes represent a small fraction of fish diversity, as the vast majority are stenohaline. Among these, freshwater fishes are challenged to maintain homeostasis in the face of wide variation in solute concentration, pH, and other chemical factors, but are less well-studied than their euryhaline counterparts [4]. Moreover, it is clear that numerous fish lineages have undergone evolutionary transitions between marine-freshwater habitats, frequently resulting in modified regulation and utilization of the proteins and genes involved in osmo- and ionoregulation [3]. Thus, although euryhaline fishes provide useful systems for understanding osmoregulatory mechanisms broadly, they may be poor models for understanding the evolutionary pathways by which stenohaline lineages invade new habitats [5] or the osmoregulatory tradeoffs that produce macroecological patterns in freshwater lineages [6].

Freshwaters of the Neotropics, which probably harbor more than a fourth of all fish diversity [7], exhibit a well-known contrast between habitats that have ion-rich “whitewater” or ion-poor “blackwater” [8]. Whitewater rivers flow over geologically young soils and are circum-neutral in pH (6-7.5), exhibiting relatively high concentrations of biologically important inorganic ions (major ions 1-10 mg/L; conductivity >50 μS/cm). Blackwater rivers generally flow over low-gradient terrain with sandy, infertile soils (podzols) and are relatively meager in free essential inorganic ions (<1 mg/L) [9] but have high concentrations of dissolved organic carbon (DOC) that produces a low pH (3-5) and low conductivity (<30 μS/cm) [10]. The ionic concentration of blackwater rivers is so low they have been described as “slightly contaminated distilled water” [8]. Despite the ionoregulatory challenges posed by blackwaters, over a thousand species are known to inhabit the Negro sub-basin of the Amazon [11], the largest blackwater drainage basin in the world (Supplemental Figure 1). Approximately two thirds of Amazon species seem to be found in only a single water type [12,13], but many closely related species show complementary distributions across water types, indicating that adaptation to water type may be an important driver of fish diversity [14]. Most research on osmoregulation in Neotropical fishes has focused on broadly or seasonally eurytopic fishes (those that occur in both water types) or contrasts among species endemic to a single water type (stenotopic). These studies have inferred that tolerance of blackwater requires greater resistance to ion loss and acidification, and that native blackwater fishes respond differently to acid challenges or ion supplementation than their whitewater or eurytopic counterparts [15,16]. Moreover, the observation that most species are not distributed across habitat types, a necessary intermediate stage for colonization of new habitats, implies that there are costs to eurytopy such that ionoregulatory strategies for blackwater versus whitewater are often evolutionarily mutually exclusive [e.g., 17]. To understand such tradeoffs, research is needed on stenotopic lineages with sub-populations adapted to different habitat types, or that vary in their degree of habitat tolerance [18]. Identifying these target species requires a robust understanding of species delimitation, population structure, and ecology, information that currently is not available for most Neotropical freshwater fishes.

One important exception are the South American tucunarés or peacock basses, large, piscivorous cichlids in the genus *Cichla*. Following analysis of extensive morphological [19] and molecular data [20,21], most *Cichla* species appear closely associated with a subset of available environmental conditions [22]. Two notable exceptions are *Cichla orinocensis* and *C. ocellaris* var. *monoculus*, the latter an evolutionary significant unit of the *C. ocellaris* species complex. Both of these meta-populations are distributed across water types in heterogeneous regions that are primarily whitewater but also punctuated by blackwater rivers (*C. orinocensis* in the Orinoco Basin, *C. oc. monoculus* in the Amazonas Basin). In addition, both occur within the Negro sub-basin of the Amazonas, a large area that is almost exclusively blackwater and connects the Amazonas and Orinoco basins together (i.e. the Casiquiare River, a Negro tributary). In both species, phylogeographic data suggest that Negro populations were derived from populations in the heterogeneous regions [23,24]. However, the Negro populations have failed to expand into neighboring heterogeneous/whitewater regions, suggesting that occupation of the blackwater habitats in the Negro sub-basin has promoted adaptations that have made these Negro fishes less tolerant of the ancestral, heterogeneous-whitewater environment.

Given the macroecological patterns exhibited by *Cichla*, these species provide an ideal system for investigating adaptation of osmoregulatory mechanisms to new physicochemical stress and associated evolutionary trade-offs. Currently, few specific details are available about the molecular and regulatory pathways involved in osmoregulation by *Cichla* in these contrasting freshwater habitats. Therefore, we employed massively parallel sequencing of the transcriptome of the primary osmoregulating organ, the gill, to identify candidate genes and genetic pathways involved in modifying ionoregulation in *Cichla oc. monoculus* for blackwater and whitewater conditions. We conducted an experiment where young fishes were exposed to water with chemistry simulating blackwater versus whitewater conditions, followed by massively parallel sequencing of mRNA from the gill filaments. This experiment was designed to identify genes that exhibit strongly different patterns of expression between these conditions while minimizing the changes in expression involved in a generalized stress response, for the purpose of generating new hypotheses regarding habitat-specific physiological strategies. Broadly, we expected that strongly and differentially expressed genes would implicate the osmoregulatory mechanisms previously identified in fishes, while also indicating how those may be modified to meet the disparate physicochemical conditions observed in the Neotropics. Our results provide an informative context for further investigating the evolutionary tradeoffs in osmoregulatory adaptation, and we discuss a number of knowledge gaps that remain to be addressed.

## Methods

We obtained 18 juvenile *Cichla oc. monoculus* from Colombia, where they had been in pond culture in local water for one or more generations. To confirm their identity and geographic origin, we sequenced the mitochondrial control region using previously published primers and conditions [25]. These fishes exhibited the most common haplotype from a *C. oc. monoculus* mtDNA clade exclusive to the western Amazon (including the Colombian Amazon), a heterogeneous but primarily whitewater region [data available upon request; see Supplemental Figure 1 and ref., 20]. These fish, approximately 5 cm at the time of the experiment, were kept in a 90 gallon (340 L) glass aquarium for 34 days prior to the exposure, with water circulation/filtration by an Eheim brand canister filter with ceramic media, daily 20% water changes, 12 hr light/dark cycle, and daily feedings of locally-collected live *Gambusia affinis* prophylactically treated for bacterial and protozoan parasites with kanamycin and metronidazole several weeks before feeding. Holding water consisted of reverse osmosis (RO) with a salt mixture added to mimic an intermediate of whitewater and blackwater habitats [9], and added to a conductivity of 30±4 μS/cm: ~2.3:1:2.3:1.7 of NaCl, K_2_SO_4_, MgSO_4_•7H_2_O, and CaCl_2_•2H_2_O by mass, respectively (from Fisher Scientific). DOC (humic acid from Sigma Aldrich) was added at a rate of 10 mg/L, assuming that this humic acid is 40% DOC [26]. The pH was adjusted to 6.2 using H_2_SO_4_ and maintained ±0.6 pH units with water changes during the holding period (Figure 1). These conditions elicited no obvious signs of stress, and the fish increased in size during the holding period. Temperature was maintained at 25±1°C throughout the experiment. Temperature and conductivity were tracked with a YSI meter (YSI Inc.), and pH was monitored using an accumet AP71 (Fisher Scientific).

**Figure 1.**
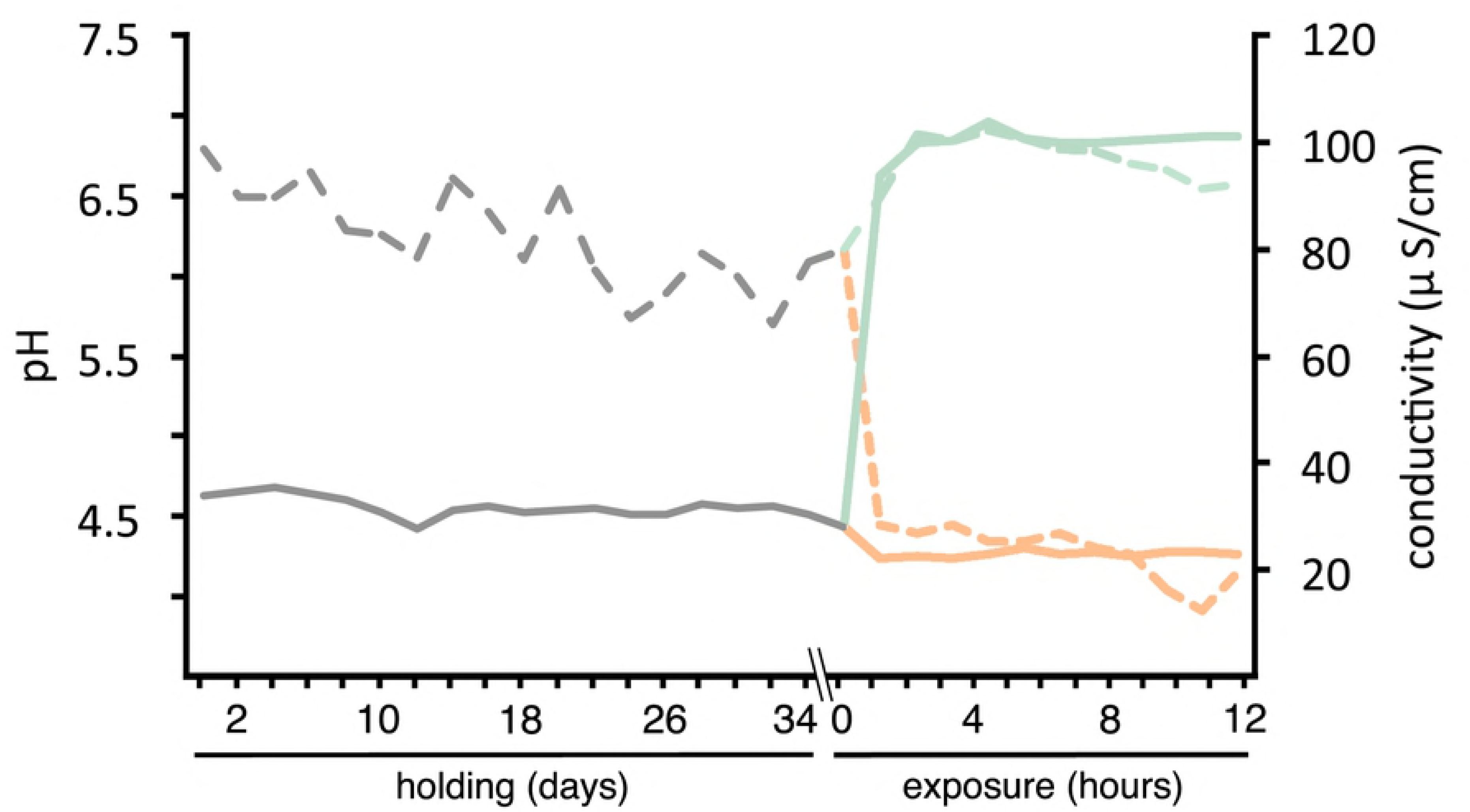
Experimental conditions for holding (days) and exposure (hours) periods. Holding period: dark gray; whitewater: green; blackwater: orange. Dashed lines: pH; solid lines: conductivity.

Fish were not fed for 48 hours prior to the exposure experiment, and a 50% water change was made 24 hours prior. Exposures were conducted over 12 hours during daylight hours on day 35 (Figure 1). Nine fish were transferred into each of two 76 L aquariums that were filled to 3 cm (~6 L) with water from the holding tank. Over the first hour, water from either the simulated “black” or “white” water was transferred slowly so that the experimental chamber was only filled at the end of the hour. All but 3 cm was removed from each chamber, and over the second hour the chambers were again filled with simulated “black” or “white” water. Simulated whitewater consisted of RO water with the same salt combination but added to a conductivity of 100±6 μS/cm, pH of 6.7±0.2, and 2 mg/L DOC. Simulated blackwater consisted of RO water saturated with DOC (between 20 and 30 mg/L) and adjusted to pH 4.2±0.3 for a final conductivity of 23±1 μS/cm. No additional salts were added to simulated blackwater, so the final chamber contained only diluted salts from the initial 3 cm of holding water [minor additional salts are also added with the Sigma Aldrich humic acid; see, 26]. Water was circulated in each chamber using a submersible pump to maintain normoxia (dissolved oxygen ≥5.5 mg/L). We acknowledge that this experimental design has no technical control, since both exposure conditions differ from holding. However, this arrangement provides for a contrast that emphasizes acclimation to novel ionic conditions in each treatment, since both experimental groups are likely to experience a degree of generalized stress response. After 12 hours, fish were euthanized by severing the spinal cord at the skull, and the gill arches excised and preserved in RNAlater (Invitrogen). Gill filaments were separated from the arches, and total RNA was extracted from the filament tissue using Trizol reagent (Invitrogen), treated with DNase (New England Biolabs), and quantified with a Nanodrop (Thermo Scientific). Total RNA was pooled equally by mass in sets of three within treatment into six total samples, and libraries were prepared with the Illumina Truseq Stranded kit (with A-tail selection) at the Texas A&M AgriLife Genomics and Bioinformatics Service. Indexed libraries were sequenced on a single lane of Illumina HiSeq 2500 with paired-end 125 bp reads. Experimental procedures were approved by the Texas A&M University Institutional Animal Care and Use Committee, 2013-0099.

A detailed description of bioinformatic and statistical procedures is available in the Supplemental Information. Briefly, *de novo* transcriptome assembly was made with several assemblers and a range of *kmer* values; various merged assemblies were also made. These assemblies were evaluated for completeness, redundancy, fragmentation, and read mapping efficacy. Quality-trimmed reads were then mapped to the optimal assembly. Even given a perfect assembly, the inherent similarity between splice variants and alleles of the same genes (hereafter, isoforms) means that mapping of reads to the correct isoform transcript can often not be done unambiguously. In addition, the analysis of differential expression (DE) of isoforms from the same gene (DIE) has less power and is more artifact-prone than length-corrected summation across isoforms from the same gene [27]. Having no reference genome, to mediate this we clustered contigs in the optimal assembly based on reads that map in common, testing for DE at a higher hierarchical level (clusters) than individual transcripts. We employed two programs that cluster transcripts based on co-mapping, Corset [28] and Rapclust [29], and with Corset, we clustered transcripts with two read co-mapping thresholds, ~30% and ~70%. These clusters were filtered to those with a sufficient expression level (length-standardized read counts per million mapped reads ≥ 1 in at least 3 samples), and tested for differential expression between treatments using three statistical packages. We employed 27 combinations of read mapping, quantification, isoform clustering, and statistical testing; to eliminate false positives, the union (overlap) of transcripts from all these combinations determined the final sufficiently expressed (hereafter SE) and differentially expressed (DE) sets. By taking as our SE and DE sets only those transcripts that were identified by the union of all methods, our bioinformatic pipeline was designed to avoid anomalies based on mapping, quantification, clustering, or statistical procedures.

The SE set (containing the DE set) was annotated using BLAST (*blastx*), with the search constrained to curated proteins (NCBI Refseq) for Nile tilapia (*Oreochromis niloticus;* hereafter *Onil*), supplemented with other cichlids (*Neolaprologus brichardi, Haplochromis burtoni, Maylandia zebra, Pundamilia nyererei*) and *Danio rerio* (hearafter “cichlid+”). Annotated transcripts were filtered to unique genes (loci) based on NCBI gene symbols, and the longest contig corresponding to each “cichlid+” gene was further BLAST against human and *Danio rerio* Refseq proteins (separately). Using the human accessions of the “cichlid+” genes, we obtained gene ontology (GO) annotations and tested for GO term enrichment using Fisher’s exact test with FDR set to 0.05 [in Blast2GOv4.1.9;, 30]. Enrichment for GO terms was tested for DE vs. SE, up-regulated vs. down-regulated, up-regulated vs. SE, and down-regulated vs. SE comparisons. We also identified which genes in the SE or DE sets were part of several well-described osmoregulatory pathways based on the gene sets annotated in the following Pathcards [31]: *prolactin signaling pathway, growth hormone receptor pathway, aquaporin mediated transport, epithelial tight junctions* (Qiagen), *tight junctions* (KEGG), and *transport of glucose and other sugars, bile salts and organic acids, metal ions and amine compounds*. Where additional genes were hypothesized to be functionally relevant for osmoregulation (see Discussion), but were not among the annotated SE set, we obtained transcripts of *Onil* from Ensembl, *tblastx* searched this against the raw transcriptome assembly, and confirmed new annotations by *blastx* search of the identified contig against Ensembl *Onil* proteins, only accepting reciprocal best matches.

## Results

It was apparent during the experimental exposure (Figure 1), following a month in common, intermediate conditions, that fishes in experimental blackwater conditions experienced greater stress than their counterparts in experimental whitewater conditions. During the second hour, fishes in the blackwater treatment moved to the corners of the aquarium and increased their ventilation rate; by the end of the experiment, a few individuals (<50%) were exhibiting a loss of equilibrium. No change in behavior from the holding period was apparent for the fishes in the whitewater treatment.

After quality trimming, the sequencing reads consisted of 190.8 million read pairs, ranging from 28.2 to 33.5 million per sample (mean 31.8). From the available *de novo* assemblies, the Transfuse [32] merger of two Binpacker [33] assemblies was considered optimal because it exhibited comparable scores and mapping rates to other top ranked assemblies but contained fewer transcripts with higher N50, implying lower redundancy and fewer mis-assemblies (Table 1, Supplemental Table 1). This transcriptome assembly of 185,480 contigs larger than 200bp (available upon request from the corresponding author), contained 281.9 Mbp and had a GC content of 44%. Mapping to these transcripts was similar across samples (92.16 to 92.71%) and mapping programs (Bowtie2 92.36%; Salmon 92.4%). Clustering transcripts based on mapping yielded between 18,203-22,997 sufficiently expressed (SE) clusters, depending on clustering algorithm and co-mapping threshold used. Principal components analyses of regularized log data from all SE clusters, with primary axes that explained 80-81% of variation, clearly separated the treatments, while the secondary axes, which explained 6-7% of variation, separated samples within treatments (Figure 2). Of the SE clusters, 7,753-9,041 were significantly differentially expressed (DE) between treatments with an FDR ≤ 0.05 (Supplemental Figure 2).

**Table 1.** Statistics and scores from Transfuse-merged *de novo* transcriptome assemblies. Top score in bold. For unmerged assembly scores, see Supplemental Table 1.

| *De novo*<br>Assembler | *kmer* Range<br>(no. *kmers*)<sup>a</sup> | Transcripts<br>>200bp<sup>b</sup> | Total Length<br>(bp)<sup>c</sup> | N50<br>Length (bp)<sup>d</sup> | Mean Contig<br>Length (bp)<sup>e</sup> | Mapped<br>Reads<sup>f</sup> | Detonate<br>Score<sup>g</sup> | TransRate<br>Score<sup>h</sup> | BUSCO<br>genes<sup>i</sup> |
| --- | --- | --- | --- | --- | --- | --- | --- | --- | --- |
| oases | 21-99 (7) | 344,662 | 2.35x10<sup>8</sup> | 835 | 682 | 1.79x10<sup>8</sup> | -4.44x10<sup>10</sup> | 0.380 | 2,552 |
| SOAPd.-Tr. | 21-99 (11) | 312,862 | 3.67x10<sup>8</sup> | 2,486 | 1,171 | 1.87x10<sup>8</sup> | -1.55x10<sup>10</sup> | 0.540 | 4,162 |
| Binpacker | 25, 32 | 185,480 | 2.82x10<sup>8</sup> | **2,648** | **1,519** | **1.87x10<sup>8</sup>** | -1.02x10<sup>10</sup> | 0.548 | 4,189 |
| Trinity | 25 | 251,811 | 2.40x10<sup>8</sup> | 1,887 | 951 | 1.85x10<sup>8</sup> | -1.17x10<sup>10</sup> | 0.542 | 4,108 |
| transABYSS | 32-96 (5) | 348,367 | 3.57x10<sup>8</sup> | 1,676 | 1,024 | 1.86x10<sup>8</sup> | **-8.66x10<sup>9</sup>** | 0.549 | 4,183 |
| ALL | *n/a* | 376,024 | 4.34x10<sup>8</sup> | 2,186 | 1,155 | **1.87x10<sup>8</sup>** | -1.87x10<sup>10</sup> | **0.562** | **4,192** |

<sup>a</sup> size range of kmers used in assembly, and where indicated, number of kmers applied
<sup>b</sup> number of transcripts larger than 200 base pairs
<sup>c</sup> combined length of assembly
<sup>d</sup> smallest contig above which 50% of the length of the assembly is found
<sup>e</sup> mean length of contigs in assembly
<sup>f</sup> total number of mapped reads (by Salmon)
<sup>g</sup> likelihood score for each assembly by Detonate program (smaller is better)
<sup>h</sup> combined score for each assembly by TransRate program (larger is better)
<sup>i</sup> number of complete conserved genes identified by BUSCO program (more is better)

**Figure 2.**
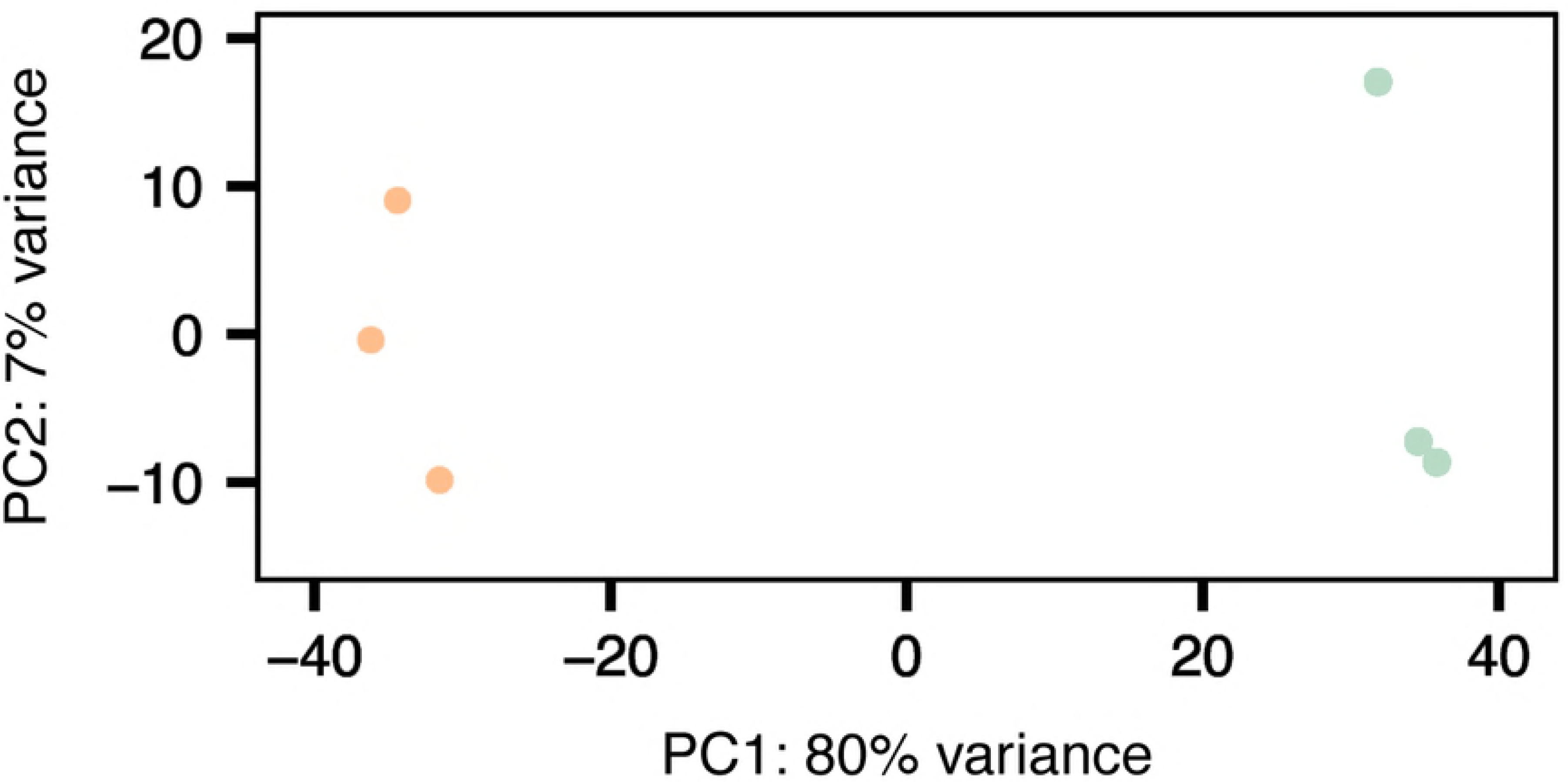
Principal components analysis of regularized log expression data from mapping and quantification with Salmon, clustering with Corset at ~30% read co-mapping. Green circles, whitewater treatment; orange circles, blackwater treatment.

The SE clusters from each mapping-quantification-clustering-testing combination contained 70,062 transcripts in common, of which 21,378 were DE in all combinations. After clustering transcripts at 98% sequence similarity with CD-hit [34], resulting in 60,567 representative transcripts, 81% (49,079) had significant hits to Refseq proteins for Nile tilapia, other cichlids, or *Danio rerio*, which corresponded to 17,379 unique protein-coding loci (hereafter, the “cichlid+” set; Supplemental Table 2). The success of blast annotation was partially correlated to the length of the contig (Pearson’s product-moment = 0.30, *P* < 2.2×10^−16^). From these “cichlid+” annotations, the DE set was found to contain 6,783 protein-coding genes, but 110 genes were found to be both up and down regulated; these were removed from the DE set (but see Discussion), which subsequently included 3,273 up- and 3,400 down-regulated genes in blackwater relative to whitewater. When BLAST against human proteins, 15,618 (90%) of the representative “cichlid+” transcripts had significant matches to 11,312 human genes (72% unique) (Supplemental Table 2). When blast against *Danio rerio* proteins, a greater percentage of representative “cichlid+” transcripts had significant hits (16,477 or 95%), which also came from a higher percentage of unique loci (13,707 or 83%) (Supplemental Table 2). The human proteins in the SE and DE sets are shown in Supplemental Table 3 along with the surveyed pathways with which they are associated. Interestingly, 242 human genes were hits for two or more “cichlid+” contigs that exhibited contrasting DE (Supplemental Table 3). Mapping of gene ontology (GO) terms for the human protein hits was successful for 15,276 of 16,477 transcripts with human annotations (98%), with a similar percentage for DE contigs (5,904 of 6,011, or 98%). Enriched GO terms are listed in Supplemental File 2. Search results for genes not identified among the SE set (and related homologs) are described below. Following common practice, below italic type refers to the gene (locus) coding for a protein (e.g. *AQP3*) and normal (Roman) type refers to the protein or protein complex (e.g. NKA).

## Discussion

### RNA-seq and Assignment of Homology

RNA-seq has tremendous potential to illuminate the ecological genomics of organisms beyond those with well-annotated genome sequences, which currently represent a tiny fraction of species and functional groups [35,36]. Meaningful results are not obtained without significant challenges, however, and our bioinformatic pipeline was designed to mediate many of the problems associated with RNA-seq data in non-model organisms [37]. Moreover, our interpretations of the biological patterns exhibited by the present data are not based on the DE or SE status of only a few genes, and so are robust to some artifacts in RNA-seq data. That being said, the interpretation of any expression pattern for both model and non-model organisms, that is, expectations of the similarity of function, regulation, etc., is dependent on the establishment of homology. In practice, this is usually based on sequence similarity to reference genes at the nucleotide or amino acid level (e.g. BLAST searches), the efficacy of which diminishes with phylogenetic distance between query and reference [38]. In addition, the presence of homologs created by duplication within the genome (paralogs) of subject or reference means these matches often show one-to-many or many-to-one relationships, obscuring the transfer of identify, function, etc. between genomes. These types of relationships are especially common between fishes and tetrapod vertebrates because of the whole genome duplication(s) early in the evolution of euteleost fishes, including cichlids [39].

Our determination of identity and function of our SE and DE transcripts was based on homology with two model organisms: Nile tilapia (*Onil*) and humans. While *Onil* is in the same family as *Cichla* and is the closest reference with a robust genome sequence and annotation available, it is still likely tens of millions of years divergent [>50;, 40]. We used the *Onil* annotations to assign our SE and DE transcripts to putative orthologous genes (one copy per haploid genome) based on amino acid similarity, a process that is subject to changes in homology (orthology/paralogy) in the intervening history between *Onil* and *Cichla*. For example, 110 out of the 6,783 *Onil* genes matched by DE transcripts were corresponded to *Cichla* transcripts in clusters that were DE in different directions. Without a full genome for *Cichla* it cannot be clarified if these result from artifacts in,clustering or DE testing, inaccurate BLAST identification, or paralogy among cichlids, and for clarity these genes were resigned to the SE set. While the percentage of genes showing this pattern is small (1.6%), in the case of paralogy, the result would mean that a greater number of unique genes (loci) are SE or DE in the present data than have been currently recognized based on assumed orthology with *Onil*.

Similarly, our identification of gene ontology and inclusion in pathways related to osmoregulation depended on homology with human proteins. While humans are significantly more distant from *Cichla* than is *Onil* (or *Danio*), the level of annotation of human proteins is unsurpassed. As with *Onil*, 242 of 6,011 DE human genes with matches to *Cichla* transcripts (4%) were found to be hits for two or more “cichlid+” genes that had contrasting DE expression (“Both” in Supplemental Table 3); a greater (unquantified) number corresponded to both “cichlid+” genes that were SE and others that were DE in a single direction. Although a small percentage of these many-to-one hits that we inspected result from erroneous BLAST identification, the vast majority appear to result from changes in homology along the human and cichlid lineages. This is substantiated by our parallel BLAST search of the “cichlid+” contigs against *Danio* which resulted in a greater percentage of unique genes compared to humans (83% vs. 72%), reflecting greater paralogy in fishes. The result is that in many cases the functional role or regulation of a human gene/protein cannot be unambiguously assigned to a single *Cichla* “cichlid+” gene. However, rather than a failing of our pipeline, we interpret this as evidence of evolutionary innovation in fishes. Following duplication, paralogs often take on functions different from the original single locus (neo- or sub-functionalization). An example of this is the prolactin receptor (PRLR) which is known to be important in euryhaline fishes for remodeling gill tissues in hyposmotic (freshwater) transition. Humans have a single *PRLR* gene which mediates the effects of prolactin on cellular signaling, but a euryhaline cichlid model for osmoregulation (*Oreochromis mossambicus;* hereafter *Omoss*) has at least two paralogs of this gene expressed in osmoregulatory tissues [41]. While *PRLR* is highly expressed in hyposmotic conditions, *PRLR2*, which contains fewer signaling domains and occurs in at least two functionally distinct isoforms, is strongly but transiently up-regulated upon *hyperosmotic* challenge, and may play a role in diminishing the effects of lingering prolactin or could have another still-unresolved role [42]. Similarly, in Ensembl *Onil* has three paralogs of *PRLR: PRLR, PRLR2*, and an unnamed third paralog (ENS0NIG00000003653), each of which corresponded to transcripts in our *Cichla* dataset that also showed hits to the single human *PRLR* (Supplemental Table 2). Moreover, we observed that *PRLR* was significantly up-regulated in blackwater relative to whitewater, *PRLR2* was significantly down-regulated, and the unnamed paralog was SE, making the human *PRLR* DE in both directions (Supplemental Table 3). Thus, while we urge caution in usage of our human annotations, even these apparent idiosyncrasies of our pipeline provide new insight into the manner in which *Cichla* responds to ionic challenge at the biochemical level. We discuss additional examples in the next section.

Finally, for simplicity, we refer to genes with significantly higher and lower expression in the blackwater treatment as “up-regulated” or “down-regulated”, respectively, although it should be noted that the opposite expression pattern (down or up in whitewater, respectively) could result in the same pattern. Similarly, because our experimental design did not include a control group (i.e. expression in holding water was not assessed), genes with increases in expression in one condition relative to holding may appear as “down-regulated” if expression was nonetheless higher in the opposing condition, while genes that exhibited expression changes in the same direction in both experimental groups relative to holding may appear in the SE set but not be identified as DE. While we expect the broad patterns of differences between conditions described here to be robust, our interpretations may be considered hypothetical even where consistent with previous results, and we look forward to future studies that test the patterns of expression in specific genes and gill cell types.

### Expression Responses to Contrasting Ionic Stress

The response to osmotic stress in fishes occurs in two phases, an acute phase and extended phase [43]. The acute phase (lasting minutes to hours) includes the activation and insertion of ion transporters and other membrane proteins in epithelial tissues as well as systemic responses like increased blood flow to osmoregulatory organs. The extended phase (lasting hours to days) includes re-modeling of osmoregulatory tissues through cell proliferation, differentiation, and selective apoptosis. These phases are activated by osmotic sensing in relevant tissues (e.g. epithelia, pituitary) and transmitted through autocrine, paracrine, endocrine, and intracellular signaling networks [42,44,45]. Our experiment, at twelve hours duration, is likely to have captured the transition between these two phases, although, by definition, it surveyed transcription-dependent responses, which are thought to make up a larger portion of the extended response. For example, among the top (ranked by difference in proportion) over-abundant specific GO terms for the DE vs. SE genes were *positive* and *negative regulation of cell proliferation, negative regulation of apoptotic process, positive regulation of cell adhesion, positive regulation of cell cycle*, *epithelial cell migration*, *positive regulation of cell activation*, and *positive regulation of apoptotic signaling pathway*, which indicates that the gill epithelia were being extensively remodeled (Supplemental File 2). Not surprisingly, these GO terms also indicated that gill epithelial remodeling was in response to external chemical stimulus, including *response to drug, response to hypoxia, cellular response to acid chemical*, and *response to toxic substance*. Changes in gill cells apparently were biased towards the cellular interface with the surroundings, given that the genes involved were enriched for the *integral component of plasma membrane, extracellular space*, and *cell surface* terms. Comparison of terms enriched in genes up-regulated vs. down-regulated (in blackwater) showed a pattern suggesting that fishes in the blackwater treatment were undergoing more transcription-based remodeling (e.g. proliferation and differentiation), especially an increase in mitochondria-rich cells that are known to carry out ion uptake/exchange. On the other hand, fishes in the whitewater treatment showed a pattern indicating a more targeted reduction in certain cell types. This was manifest by terms such as *mitochondrial translational elongation, mitochondrial translational termination, mRNA export from nucleus, protein targeting to mitochondrion, regulation of mRNA stability, gene silencing by RNA, regulation of translational initiation* being enriched in blackwater up-regulated genes, while the term *positive regulation of apoptotic signaling pathway* was enriched in genes with higher expression in the whitewater treatment.

#### Acute Phase Response

Acute phase responses are understood to be mediated by a number of peptide and steroid hormones, prostaglandins, leukotrienes, and catecholamines, some of which act in a localized fashion (auto/paracrine), while others act systemically (endocrine) [43,44]. We interrogated our DE, SE, and raw transcriptome for evidence of activity of these signaling processes and found many (Table 2), but we limit our discussion to a few. For example, the nonapeptides vasopressin and oxytocin (“vasotocin” and “isotocin” in fish, respectively) are endocrine hormones known to regulate kidney function and have antidiuretic effects in most vertebrates [43]. However, we found two paralogs of vasotocin receptor (*AVPR*) up-regulated, while two others, as well as the isotocin rector (*OXTR*), were SE, indicating that these hormones may also act to modulate salt retention in gill cells, perhaps mediated by the numerous receptor paralogs. Moreover, isotocin is also known to induce cell proliferation and differentiation in response to osmotic challenge [44]. Like vasopressin, natriuretic peptides (NPP) are known to have antidiuretic effects, but appear to more often act in paracrine fashion [43]. While *Onil* appears to have four NPP paralogs, we observed only one (*NPPC*) among our *Cichla* SE set. On the other hand, we observed up-regulation of *Cichla* transcripts matching both *Onil* copies of the NPP receptor 1 (*NPPR1*), while receptor 2 (*NPPR2*) was down-regulated (*NPPR3* was SE). Finally, endothelin, stanniocalcin, and calcitonin are known to mediate responses to acid or hyposmotic stress by regulating the activity or transcription of ion-transporters including the V-ATPase H+ pump (*ATP6V*) or epithelial calcium channels (*TRPV*) [44]. We observed transcripts for some of these signaling molecules or precursors in the SE but not the DE data sets, while one or more of the various receptor paralogs were DE, often in contrasting directions. Broadly, it appears that among signaling molecules associated with the acute phase, the effects on gill tissues may be mediated through coordinated expression of different receptor paralogs depending on the form and severity of osmotic stress.

**Table 2.** Search results for selected signaling molecules, their receptors, and accessory proteins known to moderate osmoregulation.

| Hormone | Molecule | Query/Hit<sup>a</sup> | Expression<sup>b</sup> |
| --- | --- | --- | --- |
| Cortisol |  |  |  |
|  | Glucocorticoid receptor a | 100534398 | DE down |
|  | Glucocorticoid receptor b | 100534588 | DE down |
|  | Mineralocorticoid receptor | 100712208 | DE down |
| Prolactin |  |  |  |
|  | Prolactin, long form | E-06469 | raw |
|  | Prolactin, short form | E-06467 | NP |
|  | Prolactin-like | E-14183 | raw |
|  | Prolactin receptor | 100534586 | DE up |
|  | Prolactin receptor 2 | 100534417 | DE down |
|  | Prolactin receptor-like | 100691673 | SE |
| Growth Hormone |  |  |  |
|  | Growth hormone | E-09191 | NP |
|  | Somatolactin | E-05357 | NP |
|  | GH receptor 1 (somatolactin) | 100534400 | SE |
|  | GH receptor (growth hor.) | 100534534 | DE up |
| Insulin-like growth factor |  |  |  |
|  | Insulin-like growth factor I | 100534565 | DE down |
|  | Insulin-like growth factor II | 100534429 | DE up |
|  | IGF 1 receptor | 100711993 | SE |
|  | IGF 2 receptor | 100703210 | SE |
|  | IGF-like family receptor 1 | 100700407 | DE down |
| Vasopressin |  |  |  |
|  | Vasopressin | E-15218 | raw |
|  | Vasopressin receptor 1a | 100711102 | DE up |
|  | Vasopressin receptor 1b | 100695840 | SE |
|  | Vasopressin receptor 2 | 100707194 | DE up |
|  | Vasopressin receptor 2-like | E-16572 | raw |
|  | Vasopressin receptor 2-like | E-20274 | raw |
|  | Vasopressin receptor 2-like | E-19049 | NP |
| Isotocin |  |  |  |
|  | Oxytocin | E-15235 | NP |
|  | Oxytocin receptor | E-18982 | raw |
|  | Oxytocin receptor-like | 100702815 | SE |
| Angiotensin II |  |  |  |
|  | Angiotensinogen | E-00537 | NP |
|  | Renin | E-17285 | NP |
|  | Angiotensin-converting enz. | 100712518 | DE down |
|  | Angiotensin II receptor 1 | 100702961 | DE up |
|  | Angiotensin II receptor 2 | 100702180 | DE up |

Table 2. *continued*
| Hormone | Molecule | Query/Hit<sup>a</sup> | Expression<sup>b</sup> |
| --- | --- | --- | --- |
| Urotensin II |  |  |  |
|  | Urotensin 2 | E-16900 | NP |
|  | Urotensin 2 receptor | E-19667 | raw |
|  | Urotensin 2 receptor-like | E-16919 | raw |
|  | Urotensin 2 receptor-like | E-06491 | NP |
|  | Urotensin 2 receptor-like | E-21202 | NP |
| Natriuretic Peptides |  |  |  |
|  | Atrial natriuretic peptide | E-17953 | NP |
|  | C-type natriuretic peptide | 100712558 | SE |
|  | C-type npp-like 1 | E-12915 | raw |
|  | C-type npp-like 2 | E-07496 | NP |
|  | NPP receptor 1a | 100695733 | DE up |
|  | NPP receptor 1b | 100692491 | DE up |
|  | NPP receptor 2 | 100699899 | DE down |
|  | NPP receptor 3 | 100704651 | SE |
|  | NPP receptor-like | E-13704 | NP |
|  | Corin | 100703941 | DE up |
| Vasoactive Intestinal Peptide |  |  |  |
|  | Vasoactive intestinal peptide | 101475318 | DE down |
|  | VIP receptor 1a | 100697561 | SE |
|  | VIP receptor 1b | E-05778 | NP |
|  | VIP receptor 2 | 100707237 | DE down |
| Insulin |  |  |  |
|  | Insulin | E-00343 | raw |
|  | Insulin b | E-17585 | NP |
|  | Insulin receptor a | 100697854 | DE up |
|  | Insulin receptor b | 100696191 | SE |
|  | Insulin receptor substrate 1 | 100707127 | SE |
|  | Insulin receptor substrate 2a | 100704325 | DE up |
|  | Insulin receptor substrate 2b | 100699855 | DE up |
|  | Insulin receptor substrate 4 | E-11878 | raw |
| Glucagon |  |  |  |
|  | Glucagon | E-18307 | raw |
|  | Glucagon b | E-08919 | NP |
|  | Glucagon receptor | 100700989 | DE down |
|  | Glucagon receptor b | 100692999 | SE |
|  | Glucagon-like 2 receptor | E-19565 | NP |

Table 2. *continued*
| Hormone | Molecule | Query/Hit<sup>a</sup> | Expression<sup>b</sup> |
| --- | --- | --- | --- |
| Parathyroid Hormone |  |  |  |
|  | Parathyroid hormone | E-11680 | NP |
|  | PTH receptor 1a | E-09467 | raw |
|  | PTH receptor 1b | E-11572 | NP |
|  | PTH receptor 2 | E-04451 | NP |
| Vitamin D |  |  |  |
|  | Vitamin D3 receptor A | 100696631 | DE up |
|  | Vitamin D3 receptor B | 100695813 | SE |
| Stanniocalcin |  |  |  |
|  | Stanniocalcin 1 | 100692284 | SE |
|  | Stanniocalcin 1-like | E-12623 | NP |
|  | Stanniocalcin 2a | 100709602 | SE |
|  | Stanniocalcin 2b | 106098364 | DE up |
| Calcitonin |  |  |  |
|  | Calcitonin a | E-06147 | raw |
|  | Calcitonin b | E-15534 | raw |
|  | Calcitonin receptor | 100711460 | DE up |
|  | Calcitonin receptor-like 1a | 100707330 | DE down |
|  | Calcitonin receptor-like 1b | 100696897 | SE |
|  | Calcitonin receptor-like a | E-04617 | NP |
|  | Calcitonin receptor component | 100691006 | SE |
| Endothelin |  |  |  |
|  | Endothelin 1 | 100689848 | SE |
|  | Endothelin 2 | 102080384 | SE |
|  | Endothelin 3 | E-02242 | raw |
|  | Endothelin receptor aa | 100698766 | DE down |
|  | Endothelin receptor ab | 100706592 | DE up |
|  | Endothelin receptor ba | 100704989 | DE up |
|  | Endothelin receptor bb | E-16062 | NP |
|  | Endothelin receptor-like | 100690331 | SE |
|  | Edn-converting enzyme 1 | 100690201 | SE |
|  | Edn-converting enzyme 2a | 100698809 | SE |
|  | Edn-converting enzyme 2b | 100700204 | DE down |
|  | Edn convertin enzyme-like 1 | E-10817 | raw |

<sup>a</sup> NCBI GeneID or Ensembl ID (the latter abbreviated from ENSONIG000000XXXXX).
<sup>b</sup> Present in the differentially expressed (DE) set (up, up-regulated in blackwater; down; down-regulated), sufficiently expressed (SE) set, raw unannotated transcript assembly (raw), or not present/identifiable by Blast (NP).

The effects of the acute phase signaling molecules are diverse and not fully resolved. Some act directly to activate target proteins or enhance transcription, while others activate intracellular signaling cascades with more widespread effects [42,45]. We observed evidence of several of these signaling cascades (Supplemental Tables 2 and 3). For example, calcium/calmodulin and adenylate cyclase/cAMP represent signaling networks that result in the activation and insertion of many membrane proteins involved in the osmoregulatory response, including ion transporters and tight junction proteins, and many of these signaling participants were SE or DE in the dataset. Moreover, one of the top GO terms enriched in the up-regulated vs. down-regulated comparison was *response to cAMP* (Supplemental File 1). A number of signaling cascades are also known to affect osmoregulation through activation and transcriptional regulation leading to cell proliferation, differentiation, or turnover (e.g. MAPK or PI3K-AKT), and many of the molecules in these networks were in the SE or DE sets. Moreover, among the top GO terms enriched in the DE vs. SE comparison were *ERK1 and ERK2 cascade* and *positive regulation of JNK cascade* (both MAPK signaling), while in the up-regulated vs. down-regulated comparison, up-regulated genes were enriched for *negative regulation of MAPK cascade*, whereas down-regulated genes were enriched for *Wnt signaling pathway-calcium modulating pathway, activation of JUN kinase activity*, and *non-canonical Wnt signaling pathway via JNK cascade*. The differential expression and GO term enrichment suggest that different (though not exclusive) signaling cascades were being utilized to coordinate responses to these divergent hyposmotic challenges.

#### Extended Phase Response

The canonical extended response to osmotic challenge in euryhaline fishes is understood to be mediated through the endocrine peptide hormones prolactin, growth hormone, and insulin-like growth factor I, and the glucocorticoid hormone cortisol [46]. The effects of cortisol appear to be complex and context dependent, and increased cortisol levels are associated with both hyper- and hyposmotic challenge. In *Omoss* and *Onil* cortisol has been observed to both enhance, suppress, or act independently of the effects of prolactin or growth hormone on osmoregulatory proteins [47-49]. Consistent with this, cortisol appears to have been an important factor in both of our treatments, because the GO term *response to corticosteroid* appeared in the DE vs. SE comparison that considered all DE genes (Supplemental File 2). Interestingly, both of two paralogs of the glucocorticoid receptor (*NR3C1*), as well as the mineralocorticoid receptor (*NR3C2*), all of which may bind cortisol in fish [but see, 50], were down-regulated in the blackwater treatment (Supplemental Tables 2 and 3). While there is general consensus that prolactin acts to transform the gills of euryhaline fishes for hyposmotic conditions, it is unclear how the effects of prolactin may be mediated for different types of hyposmotic challenges. In the up-regulated vs. down-regulated comparison that contrasted the two treatments, the term *cellular response to peptide hormone stimulus* was enriched in the up-regulated set. As described above, we observed two prolactin receptors (*PRLR, PRLR2*), previously implicated in hyper/hyposmotic transition in cichlids [41], to exhibit similar contrasting expression in these two hyposmotic challenges (Figure 3). In contrast, growth hormone (GH), another endocrine peptide hormone, has been suggested to promote acclimation to hypertonic environments, its effects mediated through the auto/paracrine peptide hormone insulin-like growth factor I (IGF-I). However, although GH protein and mRNA levels increase in *Omoss* following fresh to seawater transition, the molecule itself has not consistently been shown to have significant direct impacts on osmoregulatory effectors [51,52]. Moreover, at least in *Omoss*, transcription of the GH receptor (*GHR*) is usually higher following adaptation to *freshwater*, and consistent with this, we observed that *GHR* expression was higher in the blackwater treatment. By contrast, transcription of IGF-I (*IGF1*), usually acting in auto/paracrine fashion in *Omoss* in response to GH [53], was up-regulated in whitewater, whereas transcripts of IGF-II (IGF2), which also promotes cell proliferation/differentiation, were up-regulated in blackwater. No significant change was observed in IGF receptor expression (*IGF1R, IGF2R*), but we did observe contrasting DE in several IGF binding proteins (*IGFBP*) that mediate interactions of the IGFs and their shared receptor *(IGF1R)* [54]. Notably, we also observed the GO term *positive regulation of peptide hormone secretion* enriched in the DE vs SE comparison, possibly reflecting auto/paracrine signaling (e.g. IGF) in both treatments. Thus, the actions of growth hormone, if coordinated with cortisol and paracrine action of IGF-I in the gills, may be de-coupled from GHR, which instead may have been co-opted by signaling involving prolactin and IGF-II. Some studies have speculated that, in *Omoss*, GHR may relay the somatotropic signals of prolactin in hyposmotic conditions [55,56]. Conversely, like the short form of PRLR2, some isoforms of GHR may act to mitigate the effects of growth hormone in these conditions [42]. The data from *Cichla*, with contrasting expression of *PRLR, GHR*, and *IGF2* vs. *PRLR2* and *IGF1*, are consistent with either of these scenarios, and we urge caution in interpreting these results until more detail expression studies with additional controls are performed. In any case, the effects of the PRLR and GHR are understood to be transmitted through the JAK-STAT signaling pathway, as well as through the PI3K-AKT and MAPK pathways [57], and many constituents of the JAK-STAT pathway were found in the SE set, and some, like *JAK1* and *STAT3*, were DE (Supplemental Tables 2 and 3).

**Figure 3.**
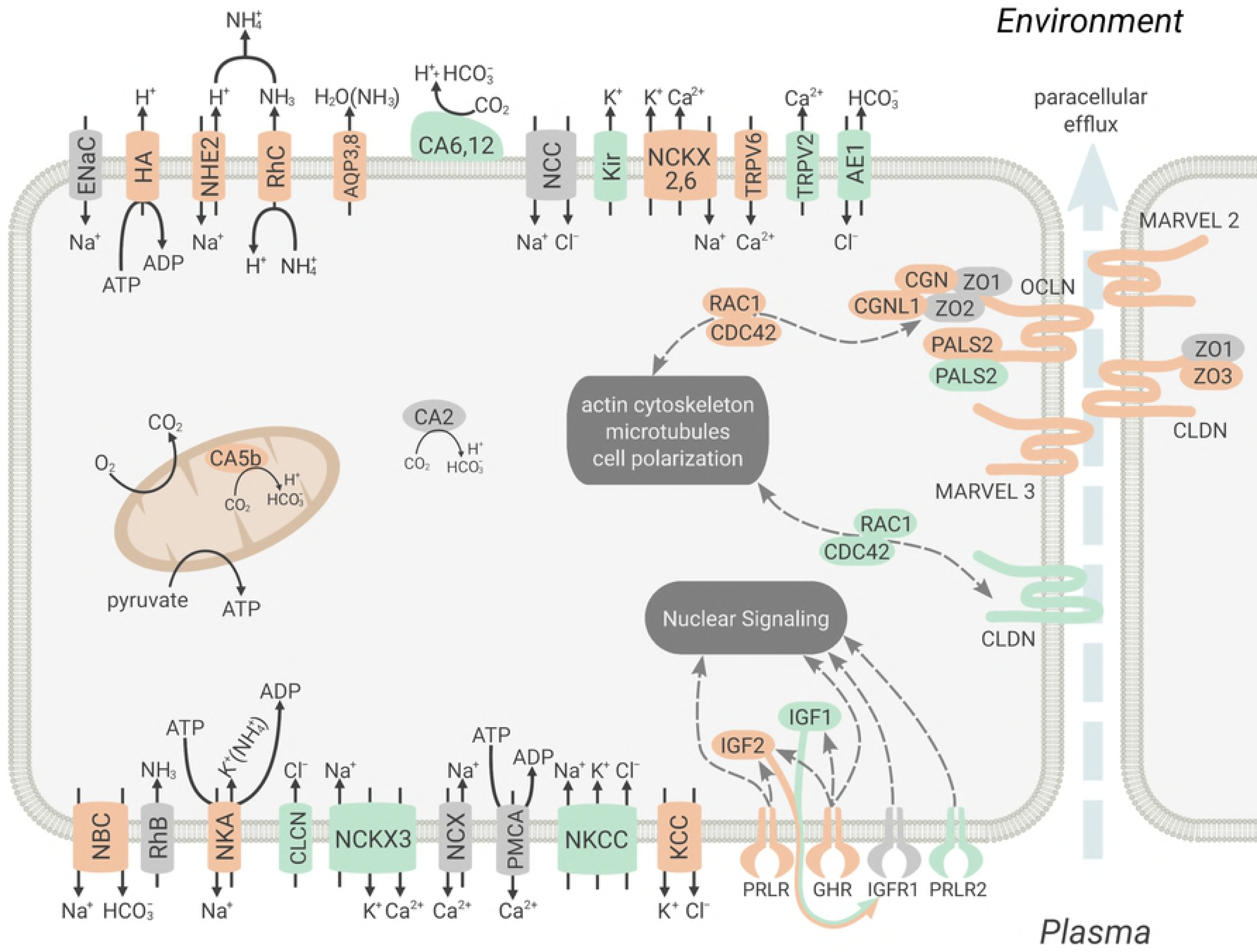
Gene expression results for select proteins, in cellular context. Color reflects expression of one or more protein isoforms (paralogs): orange, up-regulated in blackwater; green, down-regulated in blackwater; gray, no change but sufficiently expressed. Representation here is not intended to imply co-localization in the same cell type. Membrane localization and orientation reflects *Oreochromis mossambicus* (e.g. Wilson et al. 2000, Furukawa et al. 2014), but is speculative for several proteins. For protein descriptions, please see text.

The coordinated actions of prolactin, GH/IGF, and cortisol mediate paracellular permeability by regulating tight junction proteins [49,58]. In freshwater, these junctions allow little movement of ions or water (“tight”), whereas in saline conditions paracellular junctions promote the selective escape of ions down transepithelial gradients [“leaky”;, 58]. This is understood to be accomplished through increased/decreased expression of MARVEL domain-containing proteins (e.g. occludin) and selective expression of different claudin proteins; both of these groups are transmembrane proteins which obstruct and polarize the paracellular pathways as well as coordinate and communicate with gap and adherens junctions, the cytoskeleton, membrane skeleton, and signaling networks of the cell [59]. We observed contrasting DE in 25 of the 37 “cichlid+” genes identified as claudins: 14 up and 11 down (Figure 3; Supplemental Tables 2 and 3). Poor characterization of these proteins in fishes, and unclear homology with human proteins (*Onil* have ~58 *CLDN* genes; humans ~28) precludes us from predicting their effects on permeability [but see, 49]. By contrast, four of seven MARVEL domain-containing paralogs (*OCLN, MARVELD2, MARVELD3*) were up-regulated, and the others were SE, indicating that tight junctions were becoming ‘tighter’ in the blackwater treatment, and more ‘leaky’ in whitewater. It is possible that fishes adapted to whitewater may exhibit relaxed or altered mechanisms for retention of certain ions via paracellular pathways. Those losses would ultimately need to be coordinated with the transcellular transport processes discussed below.

In addition to the paracellular transmembrane proteins, a number of accessory and intermediary proteins involved in tight junction formation were also observed in the DE or SE sets. For example, the zona occludens proteins (TJP1-3), cingulin (CGN), and paracingulin (CNGL1) are known to mediate the interaction between the transmembrane proteins and the cytoskeleton [59], and several of these (*TJP3, CGN*, and *CNGL1*) were up-regulated (Figure 3; Supplemental Tables 2 and 3). Intriguingly, whereas humans have only a single copy of the small GTPases *CDC42* and *RAC1*, which respond to peptide hormone signaling and mediate tight junction assembly and cell polarity, fishes have multiple copies of each, and we observed contrasting DE among several of these. Similarly, the two paralogs of PALS2 (MPP6), a guanylate kinase that helps mediate the interaction of tight junction accessory proteins with occludin, were DE in different directions. Collectively, these observations indicate that gill remodeling in response to peptide and corticosteroid signaling involved proteins known to reduce paracellular permeability in the fish exposed to blackwater, and that evolution of this capability may have involved functional divergence of these duplicated genes.

As with tight junctions, alteration of the gill for salt uptake or secretion has been shown to be regulated by prolactin, GH/IGF, and cortisol, including in *Omoss* and *Onil* [e.g., 52]. The uptake of ions from freshwater environments and creation of transepithelial gradients is largely accomplished by transcellular transport via ion channels and pumps in coordination with passive or facilitated diffusion of metabolic solutes [3]. One of the most well documented of these is the “Na^+^/NH_4_^+^” exchange complex [60]. In this model, gradients to effectively exchange H^+^ for Na^+^ at the apical membrane are facilitated by the efflux of NH_3_ via Rhesus proteins (Figure 3). Na^+^ uptake and H^+^ excretion is hypothesized to be via Na^+^/H^+^ exchanger (NHE) or by the H^+^ pump (HA) coupled to epithelial Na^+^ channels (ENaC). As it moves across the apical gill membrane via Rhesus C (RhC), NH_3_ immediately ionizes to NH_4_^+^, reducing the external [H^+^] and partial pressure of NH_3_ (acid trapping). This process is also dependent on carbonic anhydrase (CA), which increases intracellular [H^+^] relative to the surface boundary by the hydrolysis of passively-diffusing CO_2_ to H^+^ and HCO_3_^−^, and basolateral excretion of the conjugate HCO_3_^−^ by the Na^+^-HCO_3_^−^ co-transporter (NBC), thereby also decreasing intracellular [Na^+^] relative to the boundary layer. Less clear is how NH_3_ enters the cell basolaterally, via another Rhesus protein (e.g. RhB) as NH_3_, or by active pumping through substitution of NH_4_^+^ for K^+^ in the Na^+^/K^+^ ATPase (NKA). Versions of this model are supported by experimental evidence, yet many questions remain. For example, in *Omoss* Na^+^ and Cl^−^ uptake occurs through mitochondria-rich cells (MRC) that express partially exclusive sets of ion transporters (“NHE” or “NCC” MRC), but many of these transporters move overlapping sets of ions (Na^+^, K^+^, Ca^2+^, Cl^−^, HCO_3_^−^) to energize critical gradients, and their division and coordination have not been clarified [61]. Moreover, it remains to be resolved if this Na^+^/NH_4_^+^ exchange model is effective in acidic, ion-poor environments, where the external [H^+^] and [Na^+^] are extremely unfavorable for exchange [62]. Some evidence from fish native to blackwater habitats suggests that these exchanges may be de-coupled, whereby Na^+^ uptake depends on NH_3_ excretion, but NH_3_ excretion is independent of Na^+^ uptake [15]. For example, Hirata et al. [18] found that dace native to blackwater increased somatic NH_3_ production (glutamate catalysis) to remediate Na^+^ losses, as well as NHE, CA, NBC, and aquaporin (AQP3) transcription (but not HA), whereas non-adapted dace were not able to maintain plasma [Na^+^] or pH. Similarly, when exposed to acidic, ion-poor water, *Omoss* increased both NHE and NCC (Na^+^-Cl^−^ co-transporter) transcription, but not HA [63], suggesting that Na^+^ and Cl^−^ transport may both be coordinated with NH_3_ excretion, but perhaps not via the HA/ENaC arrangement.

By contrast, *Cichla* in our acidic, ion-poor treatment exhibited up-regulation of both NHE2 (*SLC9A2*) and several subunits of HA (*ATP6V*), along with one of two paralogs of NBC (*SLC4A4*), five of seven paralogs or subunits of NKA (*ATP1*), one of two paralogs of RhC (Rhcg), and *AQP3* (Figure 3; Supplemental Tables 2 and 3). We saw no significant changes in expression (SE) in either of two paralogs of NCC (SLC12A3), NHE3 (SLC9A3), the other Rh paralogs (*Rhbg*, two paralogs of *Rhag*), or a potential ENaC (ASIC2). We did, however, note increased expression of aquaporin 8, *AQP8*, which may transport NH_3_ as well as water [64]. Interestingly, while the hydrolysis of CO_2_ invoked by the Na^+^/NH_4_^+^ exchange model is generally by cytosolic (e.g. *CA2*) or membrane-associated (e.g. *CA4*) anhydrases, expression of these did not change (SE). However, another membrane form (*CA12*) and a secreted form (*CA6*) were down-regulated, while the mitochondrial form (*CA5b*), which was shown to be critical for osmotic homeostasis in zebrafish embryos [65], was up-regulated. Although these isoforms have not been previously identified as being specifically localized or active in gill MRC, this observation is consistent with the hypothesis that MRC of fishes in the blackwater treatment were acclimating to increase intracellular [H^+^], or that in whitewater MRC were working to increase surface boundary [H^+^], to maintain gradients appropriate for acid-trapping facilitation of metabolite excretion and effective ion exchange.

Although this model focuses on Na^+^ uptake, this must also be coordinated with the transport of other ions to maintain effective electrochemical gradients. For example, it was recently discovered that *Omoss* excretes K+ in both fresh and seawater, facilitated by apical potassium channels (Kir) [61], and we observed that several paralogs of Kir (*KCNJ*) were down-regulated, consistent with increased ion conservation in blackwater (Figure 3; Supplemental Tables 2 and 3). These authors also speculated that K^+^ may be excreted through K^+^-Cl^−^ co-transporters (KCC1, KCC4), but we noted these paralogs were up-regulated (*SLC12A4, SLC12A7*), suggesting they may actually play a role in K^+^/Cl^−^ uptake or recycling. We also noted contrasting DE of several Na^+^/K^+^-Ca^2+^ exchangers, NCKX2 (*SLC24A2*), NCKX3 (*SLC24A3*), and NCKX6 (*SLC8B1*), whose roles mediating osmotic stress in fishes have not been characterized, but which generally serve to convey Na^+^ into the cell in exchange for Ca^2+^ and K^+^ [see also, 66]. Similarly, we observed contrasting DE of two epithelial Ca^2+^ channel genes (ECaC), *TRPV2* and *TRPV6*, which convey Ca^2+^ down electrochemical gradients. However, we did not observe changes in expression of either the basolateral Na^+^/Ca^2+^ exchanger, NCX (*SLC8A1*), or Ca^2+^-ATPase, PMCA (*ATP2B*), that were hypothesized to facilitate Ca^2+^ uptake in zebrafish and tilapia in coordination with ECaC [3,67]. Consistent with their role in Na^+^, K^+^, and Cl^−^ secretion in *Omoss* in seawater [68], we observed expression of one of two paralogs of the Na^+^-K^+^-Cl^−^ co-transporter, NKCC (*SLC12A2*), down-regulated in blackwater, whereas the apical chloride channel CFTR was only present in the raw transcriptome. Surprisingly, there was also down-regulation of the basolateral Cl^−^ channel (CLCN1) and the Cl^−^/HCO_3_^−^ exchanger (AE1, *SLC4A1*), both of which have been suggested to facilitate Cl^−^ uptake in freshwater *Omoss* [69,70,71; but see 59]. Finally, we identified many other DE or SE proteins known to be involved in transport of other important inorganic ions (Mg^2+^, Zn^2+^, Cu^2+^, Fe^2+/3+^) as well as amino acids and other organic ions (Supplemental Tables 2 and 3). Overall, expression of these ion transporters is largely consistent with models describing hyposmotic stress, although the specific protein isoforms and genes involved appear to vary somewhat between *Cichla*, euryhaline cichlids, and other fishes, and indicate a significant amount of functional divergence among paralogs depending on the nature and degree of the osmotic stress.

### Future Directions

We observed extensive changes in expression of genes involved in gill remodeling processes in response to ionic stress under conditions that mimicked natural environmental gradients. Our experiment contrasted conditions in whitewater and blackwater habitats, which sometimes occur in close proximity in South America and between which fish of some species disperse, though the majority of Amazonian fishes appear to be found at either end of this gradient [13]. Responding to these ionic challenges apparently involved many of the same mechanisms and regulatory pathways as fishes in hyper/hyposmotic transitions, although with some notable deviations. An RNA-seq study of an Amazonian characin, *Triportheus albus*, from rivers of different water types observed differential expression in a number of these same transporters [72]. While this study is not directly comparable because experimental subjects were surveyed *in situ*, and it is unclear whether observed expression differences result from plasticity or population-specific, constitutive expression [i.e., 14], these similarities suggest that acclimation to physicochemical challenges, and potentially also adaptation, utilize many of the same mechanisms to cope with ionic gradients of significant magnitude largely regardless of the relative tonicity of environment to body fluids [73]. In addition, our observations of several examples of contrasting expression of fish-specific paralogs of genes with known importance in human osmoregulation indicates the importance of functional diversification of these gene families in fishes for transitions among habitats with distinct physicochemistry [39]. Nevertheless, it remains unclear how many of these patterns are generalized among freshwater or Amazonian fishes.

While we observed expression of many transcripts putatively involved in osmotic stress, the system requires further study. For example, studies to date have revealed that the degree of correspondence between mRNA transcription and actual changes in protein abundance and activity can vary widely [74]. In addition, the interactive partners and cellular localization of proteins constrains their immediate function, and many of the proteins we identified are unresolved with respect to localization in the tissues or physicochemical conditions surveyed, though their correlated expression is indicative of activity in related pathways. Furthermore, it is clear that the fish gill is not a homogenous population of cells, being comprised of pavement, mucosal, neuroepithelial, and multiple types of MRCs in spatially varying proportion across the lamellae, filament, and arch epithelia [75]. However, our tissue samples consisted of homogenized filaments including blood cells, and we can directly corroborate the derivation of observed transcripts from osmoregulatory cells. Indeed, even the morphology of MRCs is not homogenous and plays an important functional role, with follicular apical crypts or pits creating ionic microenvironments at the apical membrane. This MRC morphology has been observed in cichlids inhabiting acidic, ion-poor, and hypoxic waters as well as hyperosmotic environments [63,76,77]. Our findings point to extensive molecular interactions and co-regulation in coping with ionic stress, and resolution of other details of these processes will be important for achieving a robust understanding of osmoregulatory physiology and adaptation.

Several avenues also remain to be explored in our own data. We chose not to explore differential isoform expression (DIE) because we lack splicing models for most genes; nonetheless, there are well known isoforms for some of the key osmoregulatory participants [e.g. the short form of PRLR2;, 41]. Many of the genes in the DE or SE sets likely exhibited functionally distinct isoforms whose expression may have varied between treatments, but our bioinformatic pipeline subordinated these patterns (while accounting for transcript length) to gene level expression. We also acknowledge that 19% of SE transcripts were not identifiable with the selected reference sets. A significant proportion could represent non-coding RNA that would not be identified in our search against protein databases, and indeed a casual search with some of these unidentified transcripts showed significant matches to *Onil* non-coding RNA genes (results not shown). The functional role of non-coding RNA is an area of active research and lies beyond the scope of the present study. Finally, additional genes undoubtedly exhibited DE in our experiment but were not identified due to low expression or other technical artifacts. We therefore consider our findings conservative with regard to identification of all DE genes.

Our findings provide an initial step for research exploring evolutionary tradeoffs in adaptation to novel osmotic environments and stimulate many new questions. The fish utilized here are native to a region dominated by whitewater habitats, and they were tested for responses to whitewater and blackwater conditions after being acclimated to conditions that were intermediate. Thus, our procedure captured only one of several dimensions important in adaptation to novel physicochemical environments, including population-level variation, developmental plasticity, and epigenetic effects [e.g., 5,6]. Physiological observation of Negro River fishes has indicated that some species possess higher affinity Na^+^ uptake mechanisms and decoupling of NH_3_ excretion from Na^+^ import/H^+^ export [e.g., 15]. In addition, the DOC found in Negro habitats may have unique chelating properties utilized by native fishes to minimize ion loss [78,79], and attempts to recreate this with other DOC sources (including Sigma humic acid) have produced mixed results [26,80,81]. The latter observation may partially explain why several Negro fishes tested in native water have shown lower dependency on external Ca^2+^ to charge paracellular junctions [82,83].

As a result, while acclimation to the hyposmotic gradients created here mimicked patterns seen in euryhaline fishes transitioning between seawater and freshwater, it remains to be seen if this would be true of blackwater-native *Cichla*. Based on results from our experiment, we hypothesize that *Cichla* endemic to the Negro River sub-basin appear to have been unable to colonize whitewater regions, in part, due to an inability to efficiently regulate NH3 excretion via boundary-layer acidification or to modulate the retention of some ions through paracellular or transcellular pathways [36,84]. Possible reasons for this could be isoform expression canalization, amino-acid substitutions in effector proteins (ion transporters or tight junction regulators), insensitivity of osmoregulatory complexes to the GH/IGF-I regulatory axis, or insensitivity of the axis itself to the ion concentrations common in whitewater habitats [e.g., 85,86], any of which could reflect ionoregulatory adaptation to blackwater that becomes maladaptive in whitewater. However, *Cichla oc. monoculus* is distributed across water types in the Amazon, and molecular data suggest that these populations are connected by low to moderate gene flow [20,23]. It will be important to assess if gene flow between proximal whitewater and blackwater habitats in the central Amazon constrains ionoregulatory adaptation. If there is an antagonism between adaptation and gene flow among sub-populations in different water types, this could explain why fishes in homogenous regions, like the almost exclusively blackwater Negro sub-basin, would be less tolerant of whitewater than their counterparts from heterogeneous regions: reduced gene flow-selection antagonism facilitates fixation of blackwater-adaptive alleles in the Negro [e.g., 17]. However, dispersal across habitat types in heterogeneous regions would depend first on the ability of individual fish to tolerate a range of physicochemical conditions, and though little data are available on the breath of physicochemical tolerance in *Cichla*, observations from the current experiment, in which blackwater elicited significant stress in fish from the heterogeneous, western Amazon, suggest that tolerance is not broad. However, this also highlights the unknown influence of developmental plasticity and epigenetics on osmoregulation. For example, Moorman et al. [87] observed that *Omoss* raised in tanks mimicking the temporal variation in ionic concentration of tidal habitats successfully transitioned from fresh to seawater, while those raised in freshwater-only environments could not. *Cichla* usually exhibit site fidelity, but occasionally disperse over several kilometers [88], and consequently fish that encounter environmental variation during early developmental stages may be more capable of efficient ionoregulation across habitats as adults. Considerable additional data will be needed to address these questions. The transcriptomic findings presented here provide a foundation for research addressing both proximal and ultimate mechanisms influencing biogeographic and diversification patterns in *Cichla* and other freshwater fishes.

## Acknowledgements

The authors appreciate assistance by personnel managing the Texas A&M University-Corpus Christi High Performance Computing Cluster where several bioinformatics procedures were run. We are grateful for helpful discussions with the members of the Marine Genomics Lab (TAMU-CC) and Institute for Biodiversity Science and Sustainability (CalAcademy). A.L. Val and C.M. Wood graciously provided comments on a previous version of this manuscript. Figure 3 was created with the skillful assistance of P. Dimens. This article is publication number XX of the Marine Genomics Laboratory at Texas A&M University-Corpus Christi.

## Authors’ Contributions

SCW and KOW conceived of the study, and designed the experiment with JJC. SCW and DS conducted the experiment, and GW performed laboratory procedures leading to library preparation by TAMU Agrilife Genomics. SCW performed bioinformatics processing and statistical analyses, with assistance from CMH. All authors contributed to interpretation of the results and editing the manuscript.

## Availability of Data and Materials

Raw sequence data has been deposited with the NCBI Short Read Archive as XXXXX. The transcriptome assembly used for gene expression quantification is available upon request from the corresponding author.

## Supplemental Tables

Supplemental Table 1. Statistics and scores from Transfuse-merged and constituent (pre-merge) *de novo* transcriptome assemblies. K: kmer; # transcripts (>200bp); number of transcripts in the resulting assembly larger than 200 base pairs; N50: smallest contig above which 50% of the length of the assembly is found; mean: mean length of contigs in assembly in base pairs; length (bp): combined length of assembly in base pairs; Detonate score: likelihood score for each assembly by Detonate program (smaller is better); TransRate score: combined score for each assembly by TransRate program (larger is better); TRate good: number of contigs in assembly in the “optimal” set; % good: percent of contigs in assembly in the “optimal” set; TR opt.score: score if only “good” contigs are considered (score is generated by iteratively adding high scoring contigs); total mappings: total number of mapped reads (by Salmon); % good mappings: acceptable mappings by Transrate criteria; BUSCO: number of complete conserved genes identified by BUSCO program (more is better); BUSCO p-mc: proportion of complete genes in represented by multiple transcripts; BUSCO mc: number of complete genes in represented by multiple transcripts; BUSCO miss.: number of genes from the BUSCO set that were not recovered; BUSCO out: “short” output from BUSCO program.

Supplemental Table 2. Unique Refseq protein hits to *Oreochromis niloticus*, supplemented with *Neolaprologus brichardi, Haplochromis burtoni, Maylandia zebra, Pundamilia nyererei*, and *Danio rerio* (“cichlid+”; see text), for the sufficiently expressed and differentially expressed transcript assemblies. Longest_contig: name of the longest contig blast-annotated to that gene; length(bp): length in base pairs of that contig; Expression_Change: expression change in the blackwater treatment relative to the whitewater treatment across bioinformatic combinations; Mean_Log_Expression: Mean counts per million expression across six samples from limma voom with Corset clustering at ~70% read co-mapping (-d 0.3) and quantification with Bowtie2/RSEM, on log2 scale; Log_Fold_Change: log2 fold change of blackwater relative to whitewater samples; FDR: false discovery rate, i.e. p-value adjusted for multiple tests; cichlid+_ncbi_accession: NCBI protein accession for cichlid+ annotation; cichlid+_ncbi_geneID: NCBI GI for gene corresponding to protein accession for cichlid+ annotation; cichlid+_nbci_symbol: NCBI gene symbol for gene corresponding to protein accession for cichlid+ annotation; cichlid+_description: NCBI gene description for gene corresponding to protein accession for cichlid+ annotation; human_ncbi_accession: NCBI GI for gene corresponding to protein accession for human annotation; human_ncbi_geneID: NCBI gene symbol for gene corresponding to protein accession for human annotation; human_symbol: NCBI gene symbol for gene corresponding to protein accession for human annotation; human_description: NCBI gene description for gene corresponding to protein accession for cichlid+ annotation; danio_ncbi_accession: NCBI GI for gene corresponding to protein accession for *Danio rerio* annotation; danio_ncbi_geneID: NCBI gene symbol for gene corresponding to protein accession for *Danio rerio* annotation; danio_symbol: NCBI gene symbol for gene corresponding to protein accession for *Danio rerio* annotation; danio_description: NCBI gene description for gene corresponding to protein accession for *Danio rerio* annotation.

Supplemental Table 3. Unique hits to human proteins for the sufficiently expressed and differentially expressed transcript assemblies. For differentially expressed transcripts, the expression for the blackwater treatment relative to the whitewater treatment is shown. Genes present in surveyed osmoregulatory pathways are identified: pr, *prolactin signaling pathway;* gh, *growth hormone receptor pathway;* aq, *aquaporin mediated transport*, tjc, *epithelial tight junctions* (Qiagen) or *tight junctions* (KEGG); smt, *transport of glucose and other sugars, bile salts and organic acids, metal ions and amine compounds*. For genes that are differentially expressed, the expression for the blackwater treatment relative to the whitewater treatment is indicated. * Duplicate hits show contrasting expression, most often due to many-to-one paralogy; see text.

## Supplemental Figures

Supplemental Figure 1. Map showing localities from which the same mitochondrial control region haplotype as the experimental fish was sampled (red), from which the containing mtDNA clade was sampled (orange), from which other *Cichla oc. monoculus* haplotype clades were sampled (blue), and where other evolutionary significant units of *Cichla ocellaris* were sampled (black). The Amazonas and Orinoco basins are identified, along with the Negro sub-basin, and the Casiquiare River that connects the Amazonas and Orinoco Basins. Major blackwater areas are identified by shading; smaller blackwater rivers occur sporadically throughout the lowland Amazonas and Orinoco basins.

Supplemental Figure 2. Determination of the union of differentially expressed among 27 combinations of clustering (Rapclust, Corset with −d 0.3, Corset with −d 0.7), mapping/quantification (salmon to salmon, bowtie2 to salmon, or bowtie2 to RSEM), and statistical procedure (DESeq2, EdgeR, or limma). The union of clusters is evaluated for statistical procedure and mapping/quantification, while for clustering algorithms, where cluster names are not comparable, the union of transcripts is made. Determination of sufficiently expressed transcripts was similar except filtering produced the same results regardless of statistical procedure (total nine combinations).

## Supplemental Files

Supplemental File 1. GO term enrichment. Tags: GO terms were “over”-enriched or “under” enriched; GO ID: standard ID number of GO term; GO Name: GO term description; GO Category: biological arena to which GO term refers; FDR: false discovery rate, i.e. p-value corrected for multiple testing; P-Value: uncorrected p-value; Nr Test: number in the test group annotated with GO term; Nr Reference number in the reference group annotated with GO term; Non Annot Test: number in the test group not annotated with GO term; Non Annot Reference: number in the reference group not annotated with GO term; difference in proportions: difference between the numbers in each test or reference group that were annotated with the GO term relative to those in each group that were not annotated with this term; difference in numbers: absolute difference in numbers in each test or reference group that were annotated with the GO term

